# NMDA receptor-dependent dynamics of hippocampal place cell ensembles

**DOI:** 10.1101/181131

**Authors:** Yuichiro Hayashi

**Affiliations:** Frontier Research Center for Post-genome Science and Technology, Hokkaido University, Kita-21 Nishi-11 Kita-ku, Sapporo, Japan 001-0021; Institute of Biomedical Sciences, Kansai Medical University, 2-5-1 Shin-machi, Hirakata, Osaka 573-1010, Japan

## Abstract

Place cell activity in the hippocampus constitutes a neural representation of space. The dynamics of the place cell activity for familiar environment changes gradually over time, suggesting that this temporal dynamics enables to allocate different neural codes for spatially identical but temporally different episodes. To understand the mechanisms determining the dynamics of place cell populations, activity of hippocampal CA1 neurons was imaged during repeated performance in a spatial memory task. Comparing ensemble representations among multiple task sessions revealed that overlap rate of active place cell population was time-dependent, but independent of the number of tasks within a fixed time. This time-dependent change of hippocampal ensemble activity was suppressed by the administration of an NMDA receptor antagonist. These results suggested that the gradual change of activity pattern works as a time code, and NMDA receptor-dependent processes forms the code.

---

Temporal and spatial context of events are fundamental components of episodic memory (Tulving, 2002). The hippocampus has been shown to play an important role in episodic memory formation (Allen and Fortin, 2013; Rolls, 2010; Scoville and Milner, 1957; Tulving and Markowitsch, 1998). Place cells in the hippocampus are activated at distinct locations within a given space, suggesting that these cells provide spatial context for an episodic memory (Moser et al., 2015; O’Keefe and Dostrovsky, 1971; O’Keefe and Nadel, 1978). Whereas coding for space has been extensively studied, coding for time in the hippocampus was scarcely studied until recently. Firing sequences of hippocampal neurons are observed in temporally structured tasks, which represents the flow of time from hundred milliseconds to minutes (Eichenbaum, 2014; MacDonald et al., 2011; Pastalkova et al., 2008). Another possible representation of time in the hippocampus is active cell turnover. Population activity of place cells in relation to a given environment changes dynamically over time from minutes to weeks, (Mankin et al., 2015, 2012; Manns et al., 2007; Rubin et al., 2015; Ziv et al., 2013). In particular, for timescales longer than one day, these changes are mainly the result of turnover of active neurons from session to session (Rubin et al., 2015; Ziv et al., 2013). Therefore, events occurring at different times in the same place are assigned to different neural codes in the hippocampus. They can be used to store the temporal context of episodic memory (Mankin et al., 2015, 2012; Manns et al., 2007; Rubin et al., 2015; Ziv et al., 2013). However, the factor responsible for the temporal dynamics of hippocampal activity is unknown. If the turnover of active place cells codes time, the rate of turnover will always be constant. Alternatively, when the turnover codes experience, the rate of turnover will change depending on the frequency of experience. To test these hypotheses, calcium activity of mouse hippocampal CA1 neurons was imaged during repeated performance of spatial memory tasks, and the rate of change in population activity of place cells among multiple sessions were measured. In addition, contribution of N-methyl-D-aspartate (NMDA) receptor to the place cell turnover was tested.

## Results

Place cell activity in the hippocampal CA1 region was recorded using a combination of a genetically-encoded calcium indicator and wide-field fluorescent imaging (Fig. 1A, B). The mouse was head-fixed on a spherical treadmill that allowed it to run freely (Fig. 1A) (Dombeck et al., 2007). A custom virtual reality (VR) system for rodents was used to present spatial cues (Fig. 2A, B) (Dombeck et al., 2010; Hayashi et al., 2017; Holscher et al., 2005; Youngstrom and Strowbridge, 2012). Mice were injected with an adeno-associated virus (A A V) expressing GCaMP6f into the pyramidal cell layer of the hippocampal CA1 region and then implanted with a glass window (Fig. 1B). After 2–4 weeks of recovery, mice were trained to obtain a sugar water reward at a hidden goal zone in the virtual linear track (Fig. 2B). Once the mice were proficient at the task (>10 rewards per single 10 min session), activity patterns of the hippocampal CA1 cells were repeatedly measured in the same task with varying task intervals (Fig. 2C,D and 3A). Across all recording sessions (n = 8 recording sites, 7 mice, 172–492 cells per site), activity of the mice and their reward rate did not vary (Supplementary Fig. 1). Approximately one-fourth of cells exhibited location-specific activity, and basic properties of place cell activity such as mean calcium event rate, fraction of cells having a significant place field, spatial information content, number of place fields, and place field width did not change throughout the sessions (Fig. 3B and Supplementary Fig. 2). As described in previous studies, majority of place cells were only active in a subset of sessions (Fig. 3B) (Rubin et al., 2015; Ziv et al., 2013). First, the effects of interval between sessions on the recurrence probability of place cell activity were evaluated. Recurrence probability is defined as the probability for a cell to exhibit a statistically significant place field in one session has also a place field in a subsequent session (Ziv et al., 2013).

**Figure 1.**
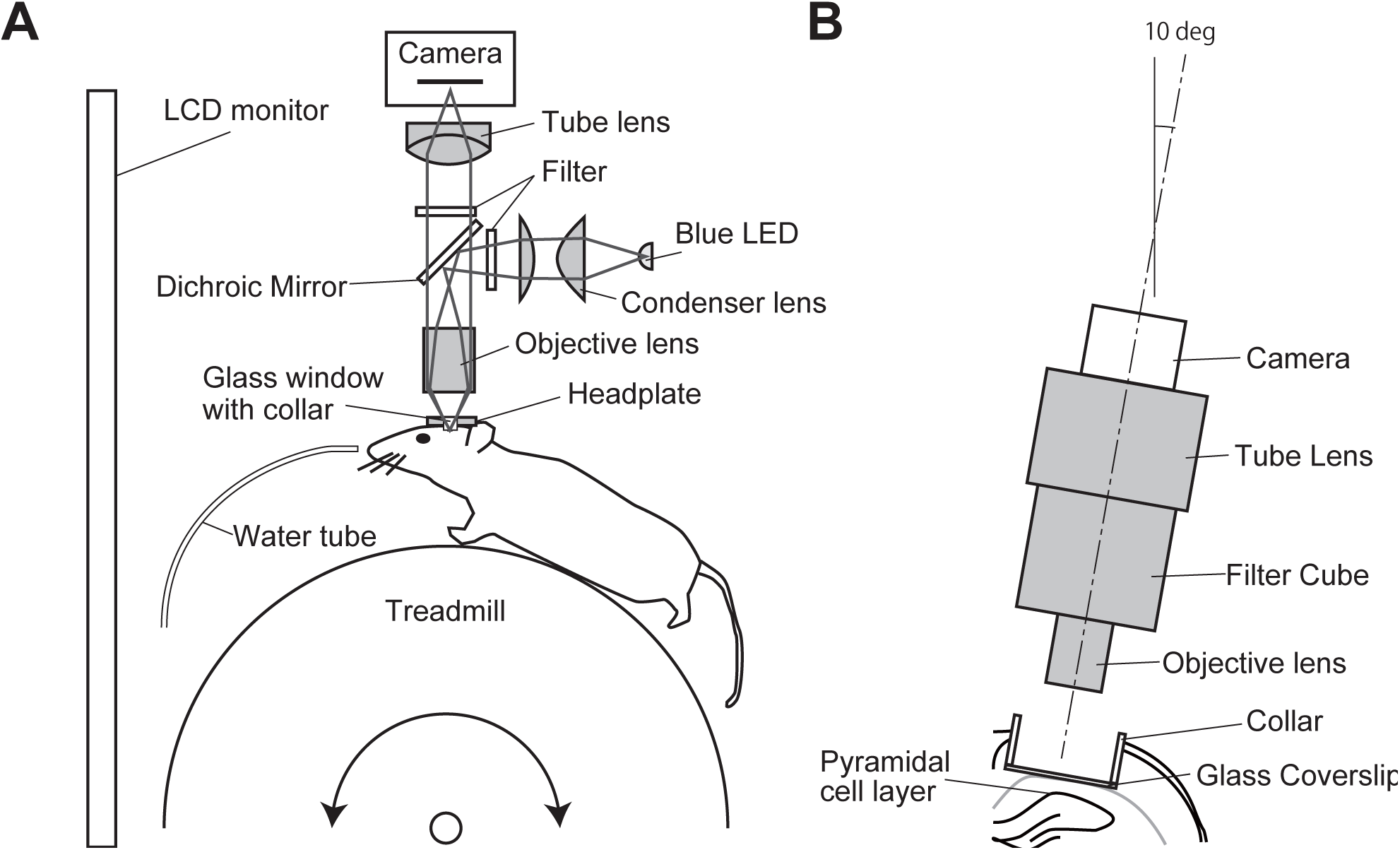
Calcium imaging of hippocampal activity in awake, head-restrained mice under VR environment. (A) Simplified schematic representation of the imaging system. The blue line indicates the illumination pathway and the green line indicates the light collection pathway. Illumination light from a blue LED was collected with a condenser lens, passed through an excitation filter, reflected off a dichroic mirror, and irradiated through an objective lens. The fluorescence image was focused on a CMOS camera. (B) Schematic representation of the experimental setup showing the chronic window implant above CA1. The optical axis of the fluorescence microscope was angled at 10 degrees. (C) View from the start point of the virtual linear track. (D) Top view of the track.

**Figure 2.**
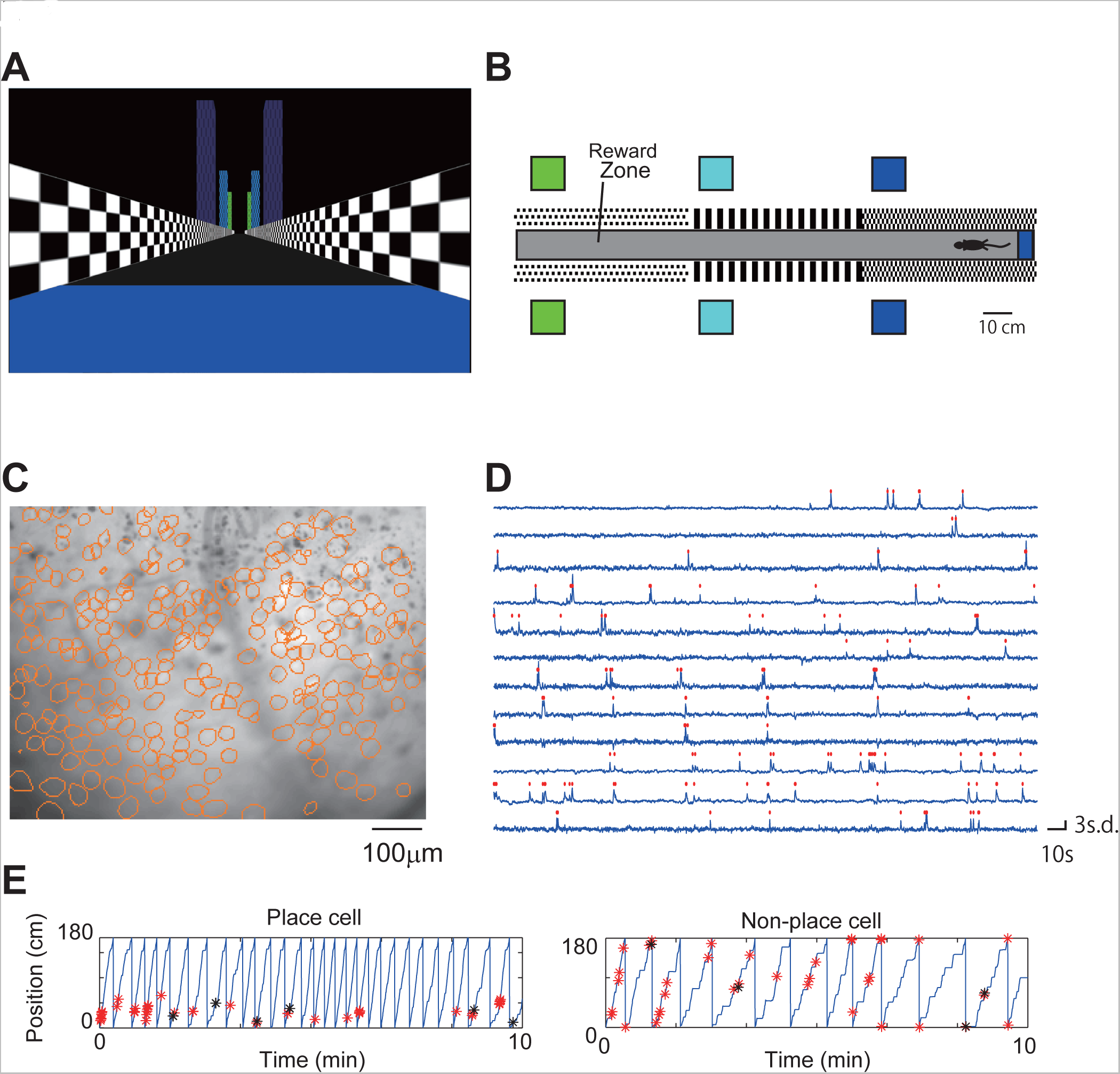
Hippocampal activity during the VR task. (A) Sample image of mean fluorescence during an imaging session. Contours of cells extracted from the fluorescent imaging movie are shown in orange circles (236 cells). (B) Relative fluorescence changes (dF/F) for 12 cells. Identified calcium transients are shown as red dots. (C) Representative CA1 cell activity. *Left*, place cell (mutual information = 1.7, Monte Carlo *P* value = 0.000); *Right*, nonspecific cell (mutual information = 0.13, Monte Carlo *P* value = 0.48). The position of the mouse along the virtual track is shown by the blue line. Identified calcium activities during running are shown with red asterisks, and those during rest are shown with black asterisks.

**Figure 3.**
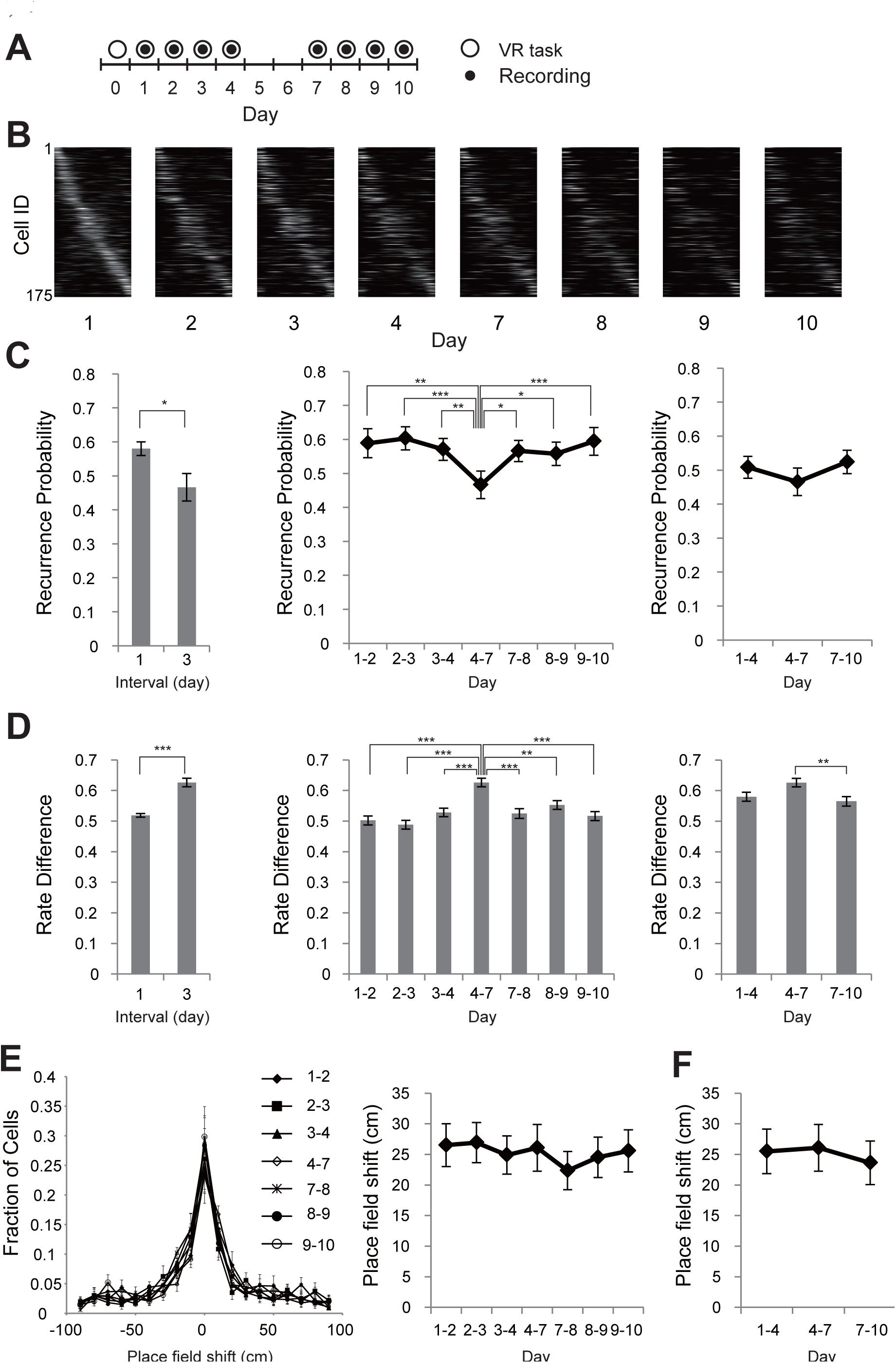
Long-term imaging of CA1 place cell activity. (A) Experimental timeline. After at least three weeks of pre-training, a series of four recording sessions was conducted on days 1–4; another series of four recording sessions was conducted on days 7–10. (B) Place field maps for the same cells on multiple days of the recording session, arranged by their centroid position on the virtual linear track. (C) If a cell had a statistically significant place field in one session, the odds that it had a place field in a subsequent session were displayed as recurrence probability. *Left*, recurrence probability for sessions with a one-day interval (recurrence between days 1 and 2, 2 and 3, 3 and 4, 7 and 8, 8 and 9, 9 and 10) versus that for sessions with a three-day interval (between days 4 and 7) were plotted. Significance was evaluated by performing an unpaired *t* test (*t* = 2.9, df = 54, *P* = 0.0044, n = 48 for 1-day interval, n = 8 for 3 day interval). *Middle*, recurrence probability for adjacent sessions were plotted. Significance was evaluated by one-way repeated measures ANOVA followed by Tukey post hoc test (*F*_6,42_ = 18.76, *P* < 0.0001, n = 8). *Right*, recurrence probability for sessions with a three-day interval. Significance was evaluated by one-way repeated measures ANOVA followed by Tukey post hoc test (*F*_2,14_ = 1.9, *P* = 0.19, n = 8). (D) *Left*, changes in the peak transient rates (rate difference) between sessions with a one-day interval versus that for sessions with a three-day interval. Significance was evaluated by performing an Mann-Whitney *U* test (*U*= 1782000, *P* < 0.0001, n = 4877 for 1-day interval, n = 856 for 3 day interval). *Middle*, rate difference for adjacent sessions. Significance was evaluated by Kruskal-Wallis test followed by Dunn’s post hoc test (*H* = 62.81, *P* < 0.0001, n = 895-702). *Right*, rate difference for sessions with a three-day interval. Significance was evaluated by Kruskal-Wallis test followed by Dunn’s post hoc test (*H* = 9.797, *P* = 0.0075, n = 856-702). (E) *Left*, Distribution of centroid shifts of place fields between adjacent recording sessions (1-2, 2-3, 3-4, 4-7, 7-8, 8-9, and 9-10). *Right*, centroid shifts of the place fields between adjacent recording sessions. Significance was evaluated by one-way repeated measures ANOVA (*F*_6,42_ = 1.622, *P* = 0.17, n = 8). (F) Centroid shifts between sessions with a three-day interval. Significance was evaluated by one-way repeated measures ANOVA (*F*_2,14_ = 0.63, *P* = 0.55, n = 8). Data are shown as mean ± SEM. Asterisks indicate significantly different (\**P* < 0.05, \*\**P* < 0.01, \*\*\**P* < 0.001; otherwise not significant).

Higher probability of recurrence was observed between 1-day interval sessions than 3-day interval sessions (Fig. 3C, *Left;* 0.58 ± 0.020, n = 48 for 1-day interval; 0.47 ± 0.040, n = 8 for 3-day interval; *P* = 0.0044, unpaired *t* test). Analysis of time-series data also indicated that recurrence probability was higher between 1-day interval sessions than 3-day interval sessions (Fig. 3C, *Middle; P* < 0.0001, n = 8, one-way repeated measures ANOVA followed by Tukey post hoc test). Then, the effects of task frequency per fixed time on recurrence probability were tested. In the first trial block, mice performed the task every day between the recording sessions (Days 1–4 in Fig. 3A). In the second block, mice did not perform the task between the recording sessions (Days 4–7 in Fig. 3A). In the last block, mice performed the task every day again between the recording sessions (Days 7–10 in Fig. 3A). No significant difference was observed in recurrence probability among these three trial blocks, indicating that task frequency had no detectable effect on the speed of place cell turnover (Fig. 3C *Right; P* = 0.19, n = 8, one-way repeated measures ANOVA). Change of place cell activity between sessions was also assayed by comparing the peak calcium transient rate in the place field. For this purpose, normalized change in calcium transient rate (rate difference: unsigned rate difference divided by rate sum) was used (Leutgeb et al., 2005). The rate difference was higher between 3-day interval sessions than 1-day interval sessions (Fig. 3D, *Left;* 0.52 ± 0.0060, n = 4877 for 1-day interval; 0.63 ± 0.014, n = 856 for 3-day interval; *P* < 0.0001, Mann-Whitney *U* test). Time-series data also indicated that higher rate difference between 3-day interval sessions (Fig. 3D, *Middle; P* < 0.0001, n = 895-702, Kruskal-Wallis test followed by Dunn’s post hoc test). Comparison of rate difference in different task frequency resulted a significant but little difference between sessions (Fig. 3D, *Right;* 0.58 ± 0.015, n = 809 for days 1-4; 0.63 ± 0.014, n = 856 for days 4-7; 0.57 ± 0.016, n = 702 for days 7-10; *P* = 0.0075, Mann-Whitney *U* test). These results were consistent with higher place cell turnover ratio in 3-day interval sessions compared with 1-day interval sessions, and also with no detectable effect of task frequency on the speed of place cell turnover (Fig. 3C).

Next, the effects of time and task frequency on the day-to-day fluctuation of place field location were evaluated. There was no significant difference in place field shift when the interval or task frequency was varied (Fig. 3E; *P* = 0.17, n = 8, one-way repeated measures ANOVA and Fig. 3F; *P* = 0.55, n = 8, one-way repeated measures ANOVA). These results are consistent with previous observations indicating that the place field location of each cell is stable over time (Lever et al., 2002; Rubin et al., 2015; Thompson and Best, 1990; Ziv et al., 2013).

Mice are nocturnal animal, sleeping during daytime and active during nighttime. If activity affects place cell turnover, the turnover rate will be different at day and night. To test this, day-night variation of place cell turnover rate was measured. Mice were housed under 12-hour light/12-hour dark photoperiod, and place cell activity was measured at start of light period (Zeitgeber time, ZT1) and at start of dark period (ZT13) (Fig. 4A). There was no significant difference in turnover rate of place cell (Fig. 4B; *P* = 0.68, n = 20; one-way repeated measures ANOVA) and rate difference of place cell activity (Fig. 4C; *P* = 0.54, n = 1371-1517 cells; Kruskal-Wallis test) beteween the daytime (between ZT1 and 13) and nighttime (between ZT13 and 1). The amount of place field shift between daytime and nighttime sessions also exhibited no significant difference (Fig. 4D; *P* = 0.32, n = 20, Friedman test). Across all recording sessions, there was no significant difference in either animal behavior or place cell properties between daytime (ZT1) and nighttime (ZT13) sessions (Supplementary Fig. 3 and 4) (n = 20 recording sites, 15 mice, 161–374 cells per site).

**Figure 4.**
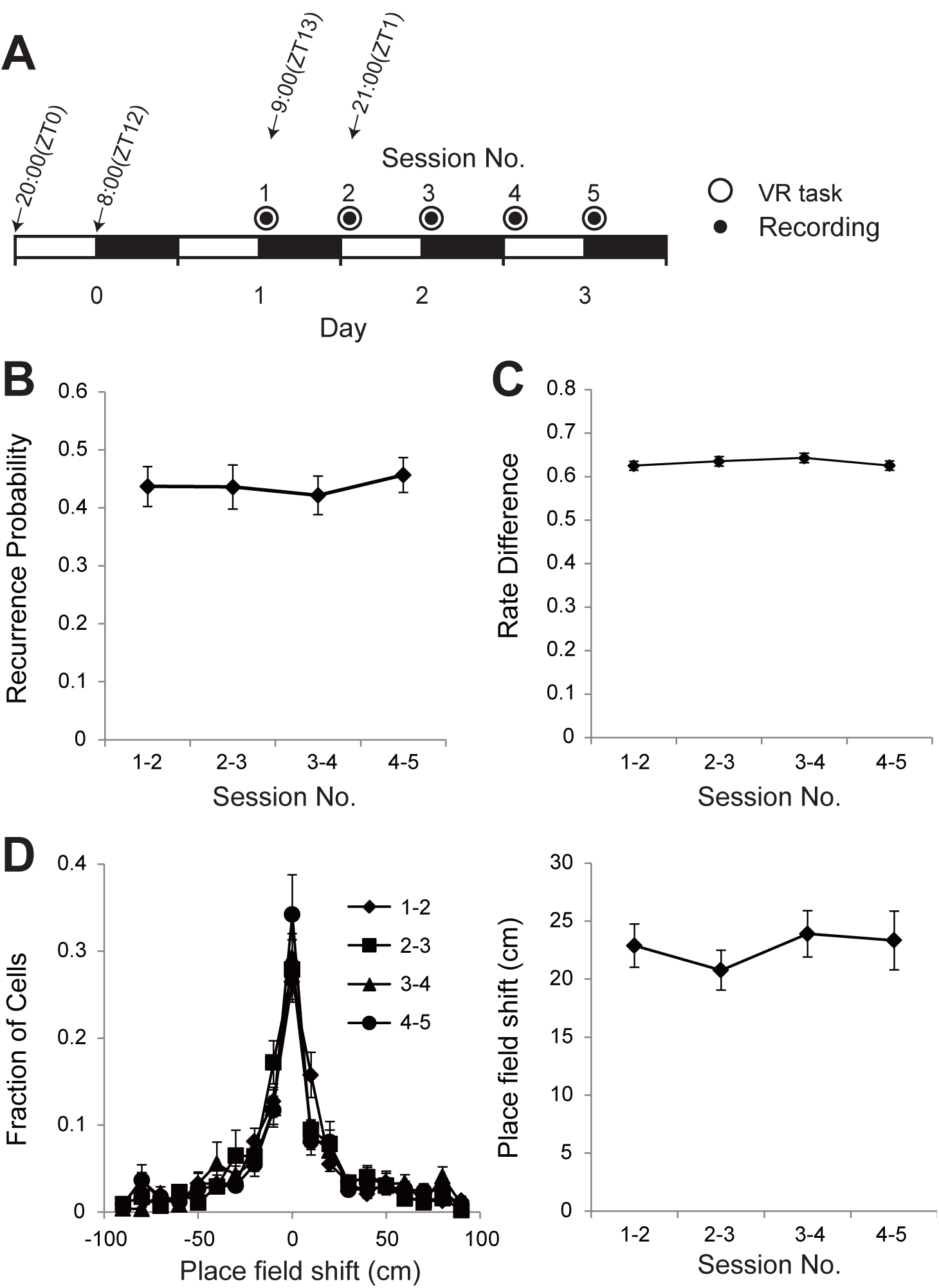
Day-night variation of place cell turnover. (A) Experimental timeline. After the pre-training period, five recording sessions were conducted bi-daily. (B) Recurrence probability for the two sessions under saline and CPP conditions. Significance was evaluated by one-way repeated measures ANOVA (*F*_4,76_ = 0.50, *P* = 0.68, n = 20). (C) Rate Difference between adjacent recording sessions. Significance was evaluated by Kruskal-Wallis test (*H* = 2.16, *P* = 0.54, n = 1517-1371). (D) Centroid shifts of place fields between adjacent recording sessions. *Left*, distributions of centroid shifts. *Right*, average centroid shifts. Significance was evaluated by Friedman test (*Q* = 2.48, *P* = 0.32, n = 20). Data are shown as mean ± SEM.

NMDA receptors have been identified to play a critical role in synaptic plasticity as well as in hippocampal memory (Malenka and Nicoll, 1993; Nakazawa et al., 2004; Steele and Morris, 1999). Previous studies have shown that inhibition of NMDA receptors sustain long-term potentiation (LTP) at perforant path-dentate gyrus synapses and promotes retention of hippocampal memory (Sachser et al., 2016; Shinohara and Hata, 2014; Villarreal et al., 2002). These studies have led to the idea that place cell turnover is also an NMDA receptor-dependent process.

To test this hypothesis, the NMDA receptor antagonist 3-((R)-2-Carboxypiperazin-4-yl)-propyl-1-phosphonic acid (CPP) was systemically administrated just after the first recording session and daily thereafter for two days (Fig. 5A). The drug administration had no effect on either animal behavior (Supplementary Fig. 5) or place cell properties including mean calcium event rate, fraction of cells having a significant place field, spatial information content, number of place fields, and place field width (Supplementary Fig. 6; n = 5 mice, 207–294 cells per animal for saline; n = 5 mice, 101–263 cells per animal for CPP). Recurrence probability of place cell activity for the 3-day interval was significantly increased in the CPP group compared with the saline group (Fig. 5B; 0.48 ± 0.053, n = 5 for saline; 0.68 ± 0.056, n = 5 for CPP, *P* = 0.032, Mann–Whitney *U* test). Rate difference in the CPP group was lower than that in the saline group (Fig. 5C; 0.59 ± 0.019, n = 514 for saline; 0.47 ± 0.021, n = 344 for CPP, *P* = 0.0005, Mann–Whitney *U* test). These results showed that NMDA receptor antagonism interfered with place cell turnover. At the same time, no significant difference was observed in the amount of place field shift between the saline and CPP groups (Fig. 5D, 22.3 ± 1.5, n = 5 for saline; 20.2 ± 1.5, n = 5 for CPP, *P* = 1.0, Mann–Whitney *U* test).

**Figure 5.**
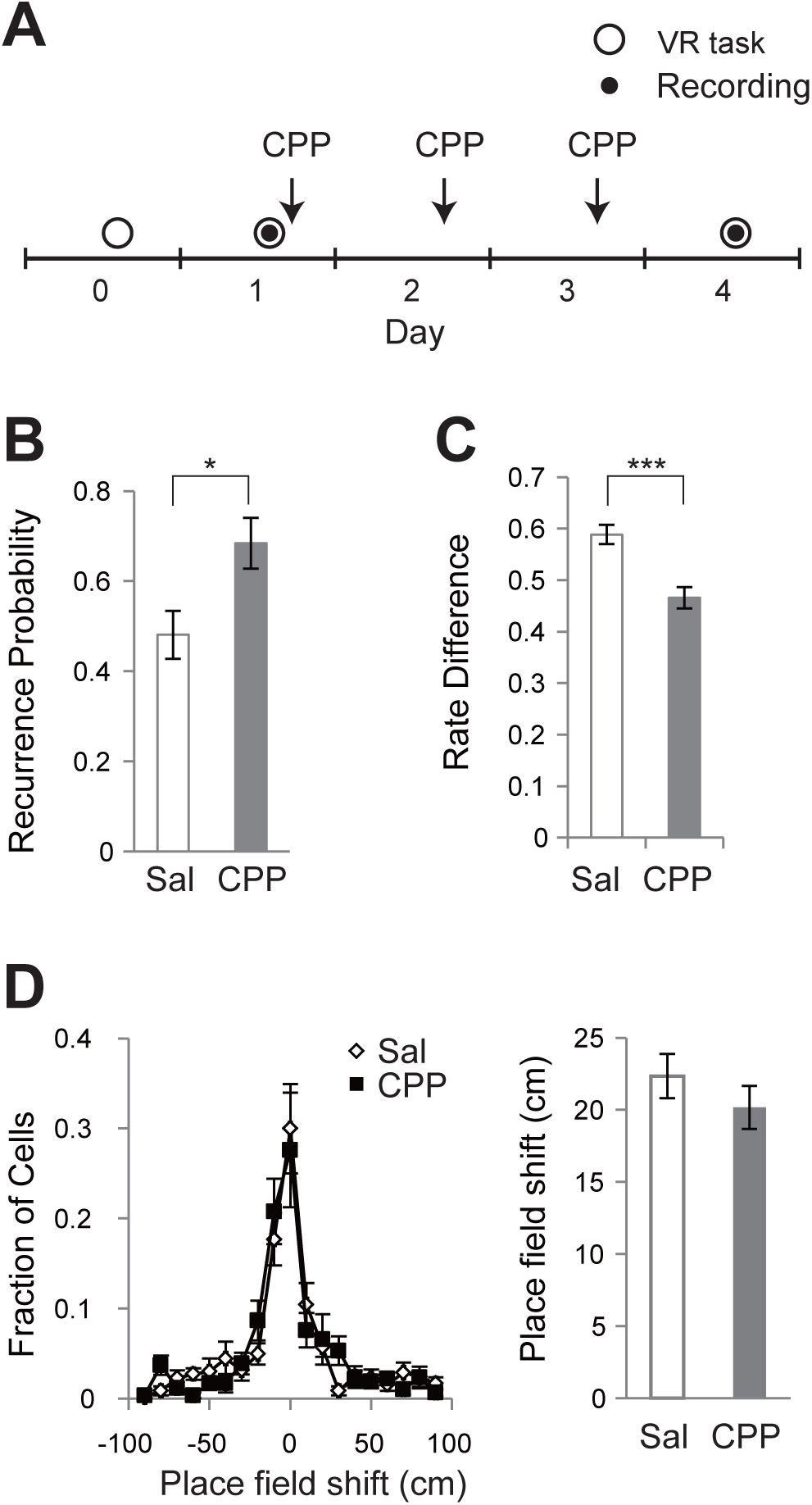
NMDA receptor antagonism reduced turnover of place cells. (A) Experimental timeline. After the pre-training period, two recording sessions were conducted on days 1 and 4. CPP was intraperitoneally administrated after the first recording session and daily thereafter for two days. (B) Recurrence probability for the two sessions under saline and CPP conditions. Significance was evaluated by Mann–Whitney *U* test (*U* = 2, *P* = 0.032, n = 5 for saline, n = 5 for CPP). (C) Rate Difference for the two sessions under saline and CPP conditions. Significance was evaluated by Mann–Whitney *U* test (*U* = 76390, *P* = 0.0005, n = 514 for saline, n = 344 for CPP). (D) Centroid shifts of place fields between the two sessions under saline and CPP conditions. *Left*, distributions of centroid shifts. *Right*, average centroid shifts. Significance was evaluated by Wilcoxon signed rank test (*W* = 1, *P* = 1.0, n = 5 for saline, n = 5 for CPP). Data are shown as mean ± SEM. Asterisks indicate significant differences (\**P* < 0.05, \*\*\**P* < 0.001; otherwise not significant).

## Discussion

Previous studies have shown that active population of CA1 place cells change dynamically over time when the animals are repeatedly exposed to a familiar environment (Rubin et al., 2015; Ziv et al., 2013). However, it was still unknown whether the speed of turnover is constant or can be modulated by task experience. To answer this question, the place cell activity in the hippocampal CA1 area was repeatedly recorded at variable task frequency. Long-term tracking of neural activity revealed that the rate of change in population activity of place cells was constant regardless of the task frequency (Fig. 3C). Moreover, no within-day variation was also found in the turnover of place cells (Fig. 4B). These results suggested that the turnover of active population of place cells can be used as a time code for memories of multiple events (Ziv et al., 2013).

A recent study suggested that this time code not only expresses the temporal relationship of events occurring at the same place but also the temporal relationship of events occurring at different places (Cai et al., 2016). The study found that overlap between hippocampal CA1 cell ensembles activated by two distinct contexts is greater when the animals are experienced within a day than when separated by longer intervals. This suggests that the place cell turnover in CA1 can work as a general time code for memories of multiple episodes.

The present study also investigated the neural mechanisms of the place cell turnover. Hippocampal memory is transient and lasts about 3–4 weeks in mice (Frankland and Bontempi, 2005). This impermanence of hippocampal memory is consistent with the high rate of spine turnover in the hippocampus (Attardo et al., 2015). Antagonists of NMDA receptor which is an important molecule regulating synaptic plasticity blocks the decay of LT P in the perforant path-dentate gyrus synapses and enhances spatial memory retention (Sachser et al., 2016; Shinohara and Hata, 2014; Villarreal et al., 2002). These studies led to an idea that the place cell turnover is also an NMDA receptor-dependent process. In the present study, NMDA receptor blockade CPP reduced the speed of place cell turnover (Fig. 5A, B), indicating that NMDA receptors play an important role in the turnover of CA1 neural ensembles. Thus NMDA receptor-dependent turnover of place cell ensembles may be one of the mechanisms determining the lifetime of hippocampal memory.

In this study, the rate of place cell turnover was constant over time irrespective of task frequency (Fig. 3). However, it cannot be denied that there are factors affecting the place cell turnover rate. One possible candidate is adult neurogenesis in the hippocampus. New neurons are continuously added to neural circuits in the dentate gyrus of the hippocampus (Zhao et al., 2008), and this process has been shown to play a significant role in memory retention. For example, forgetting is associated with increased neurogenesis and mitigated by impaired neurogenesis (Akers et al., 2014). The rate of neurogenesis can be modulated by running, environmental enrichment, or stress (Gould et al., 1997; Kempermann et al., 1997; van Praag et al., 1999), and therefore these treatments may alter the place cell turnover rate.

Although active population of CA1 place cells dynamically changes over time, their place fields show only a small fluctuation for days to weeks (Rubin et al., 2015; Ziv et al., 2013). The present study demonstrated that the fluctuation of the place field is not affected by elapsed time (Fig. 3F, *Left*), task frequency (Fig. 3F, *Right*), and NMDA receptor blockade (Fig. 5D). These results suggest the existence of a mechanism that stabilizes the place field against the continuous synaptic modifications in the brain.

## Materials and Methods

### Calcium imaging with a custom wide-field microscope

A custom wide-field microscope for in vivo calcium imaging has been previously described (Hayashi et al., 2017) (Fig. 1A). Excitation light was emitted from a blue LED (LXK2-PB14-P00; Lumileds, Aachen, Germany). The light passed through an excitation filter (FF480/40-25; Semrock, Rochester, NY) and then reflected by a dichroic mirror (FF506-Di02-25x36; Semrock) onto the tissue through an objective lens (LUCPlan FLN20/0.45; Olympus, Tokyo, Japan). Fluorescent emissions collected by the objective lens were passed through the dichroic mirror and an emission filter (FF535/50-25; Semrock). The fluorescence image, focused by a tube lens (Nikkor 50 mm f/1.8D; Nikon, Tokyo, Japan), was captured by a CMOS camera (FL3-U3-13S2M-CS; FLIR Systems, Willsonville, OR). The optical axis of the microscope was angled at 10 degrees to be perpendicular to the pyramidal cell layer of the CA1 hippocampal region (Fig. 1B).

### Animals

All animal care and use was in accordance with the protocols approved by Hokkaido University institutional animal care and use committee (project license number: 16-0042). Adult male C57BL/6J mice (10–16 weeks old) were maintained on a 12-h light/12-h dark schedule with lights of at 8:00 AM. Behavioral tasks and recording sessions occurred in the dark phase (9:00 AM-3:00 PM). In bi-daily recording experiment (Fig. 4), recording sessions started at 9:00AM (ZT13) or 9:00PM (ZT1).

### Surgery

Animals were anesthetized using isoflurane and then injected 0.1 mg/kg buprenorphine. The skull was exposed, and a small hole (<0.5 mm) was made over the right hemisphere (1.5 mm lateral to the midline, 2.3 mm posterior to the bregma). AAV1-syn-GCaMP6f-WPRE (University of Pennsylvania vector core) was diluted to 5 × 10^12^ particles/mL in phosphate-buffered saline, and 150 nL was injected into CA1 (1.2 mm ventral from the brain surface). One week after the viral injection, the animals were anesthetized and a 2.8 mm diameter craniotomy was performed. The dura was removed, and the underlying cortex was aspirated. A stainless-steel cannula (2.76 mm outer diameter, 2.40 mm inner diameter, 1.5 mm height) covered by a glass coverslip (0.12 mm thickness) was inserted over the dorsal CA1. A titanium head plate (25 mm × 10 mm, 1 mm thickness) and the skull were glued with dental cement (Shofu, Kyoto, Japan).

### Virtual reality system

The VR system has been previously described (Hayashi et al., 2017) (Fig. 1A, 2A and B). Mice ran along a virtual linear track (1.8 m long) and received a small sugar water reward (4 μl) in an unmarked reward zone (1.5 m from the start point of the track) (Fig. 2B). Once the mice reached the end of the track, they were teleported back to the start of the track.

### Behavioral training

At least two weeks after the cranial window implantation, the mice underwent water restriction (2 mL per day), and training in the virtual linear track (10- or 15-min session per day, five days per week) began. After 3–4 weeks of training, mice achieving high performance levels (>10 rewards per single 10 min session) were used for the recording session.

### Pharmacology

Mice received intraperitoneal injection of either a saline or 10 mg/kg CPP (Tocris, Bristol, UK) after the first recording session and daily thereafter for two days (Fig. 5A).

### Imaging session

The excitation light intensity was approximately 0.04 mW/mm^2^. Images were captured using FlyCapture2 software (FLIR Systems) at 10 Hz for 10 min. Calcium activity was recorded from19 imaging sites in 16 mice for the long-term imaging experiment (Fig. 3), from 25 imaging sites in 17 mice for the bi-daily experiment (Fig. 4) or from 13 imaging sites in 13 mice for the CPP experiment (Fig. 5). Recordings with a low image quality (blurry image or inconsistent focal plane throughout the experiment) were discarded.

### Image processing

ImageJ1.49q (National Institute of Health) and MATLAB 2013a (Mathworks, Natick, MA) were used for all analyses. Fluorescent images were downsampled by a factor of 4. Then lateral displacements of the brain were corrected by image translation using TurboReg (Thévenaz et al., 1998). An averaged image of the entire frame was used as a reference for image alignment. Spatial filters corresponding to individual cells were identified using a PCA-and ICA-based algorithm (Cellsort 1.0) (Mukamel et al., 2009). To select spatial filters that follow a typical cell structure, contour of half maximum intensity was calculated, and filters whose area of contour was smaller than 20 pixels or larger than 300 pixels or whose circularity of contour was smaller than 0.7 were discarded.

### Detection of Calcium transients

Calcium activity was determined by applying the spatial filters to the time-lapse image. The Calcium transients were identified by searching each trace for the local maxima that had a peak amplitude of more than 3× standard deviation from the baseline (defined as the average of the trace calculated across the entire sessions) (Fig. 2D).

### Registration of cells across sessions

The threshold spatial filters of all cells from each session were mapped onto a single image. Subsequently, the other filters were aligned to the first day’s image via affine transform using TurboReg. Following the alignment across all sessions, cell pairs with a centroid distance <7.8 μm were registered as the same neuron.

### Place fields

To analyze place fields, calcium events occurring when the animal’s velocity was less than 1 cm/s were filtered out to eliminate nonspecific activity at rest. The number of calcium transients in each bin was divided by the total occupancy time of the mouse in the bin, and then a Gaussian smoothing filter was applied (σ = 2 cm). The number of bins in which a place field had a value greater than 50% of its maximum calcium event rate determined its width. The position of the place field was defined as its centroid.

For each cell’s place field, the mutual information between calcium transients and the mouse’s location was calculated (Skaggs et al., 1993). To assess the significance of spatial selectivity, Monte Carlo *P* values were calculated. A total of 1,000 distinct shuffles of the calcium transient times were performed, and the mutual information for each shuffle was calculated. The *P* value was defined as the fraction of the shuffles that exceeded the mutual information of the cell. Cells with *P* < 0.05 were considered place cells.

Rate difference in the place field was obtained by calculating the unsigned rate difference between the peak calcium transient rates in the two recording sessions and dividing the difference by the sum of the two rates. If a cell did not have statistically significant place field in the second session, the rate was to be zero.

### Statistical analysis

Statistical analysis was performed using GraphPad Prism 5 (GraphPad Software, La Jolla, CA) and MATLAB 2013a. Data were first tested for normality with Shapiro–Wilk test and for homogeneity of variance with Bartlett’s test or the *F* test.

Normally distributed data with equal variance were compared with paired/unpaired Student’s *t* test or one-way repeated measures ANOVA followed by Tukey post hoc comparisons, as stated in the figure legends. If the data did not meet the assumption of parametric tests, non-parametric tests (Mann–Whitney *U* test, Wilcoxon signed-rank test, Kruskal-Wallis, or Friedman test) were used. Statistical significance was set at *P* < 0.05 for all statistical analyses.

## Acknowledgments

This work is supported by JSPS KAKENHI grant number 17K19436. The author would like to thank Enago (http://www.enago.jp) for the English language review.

## Figure Legends

**Fig. S1.**
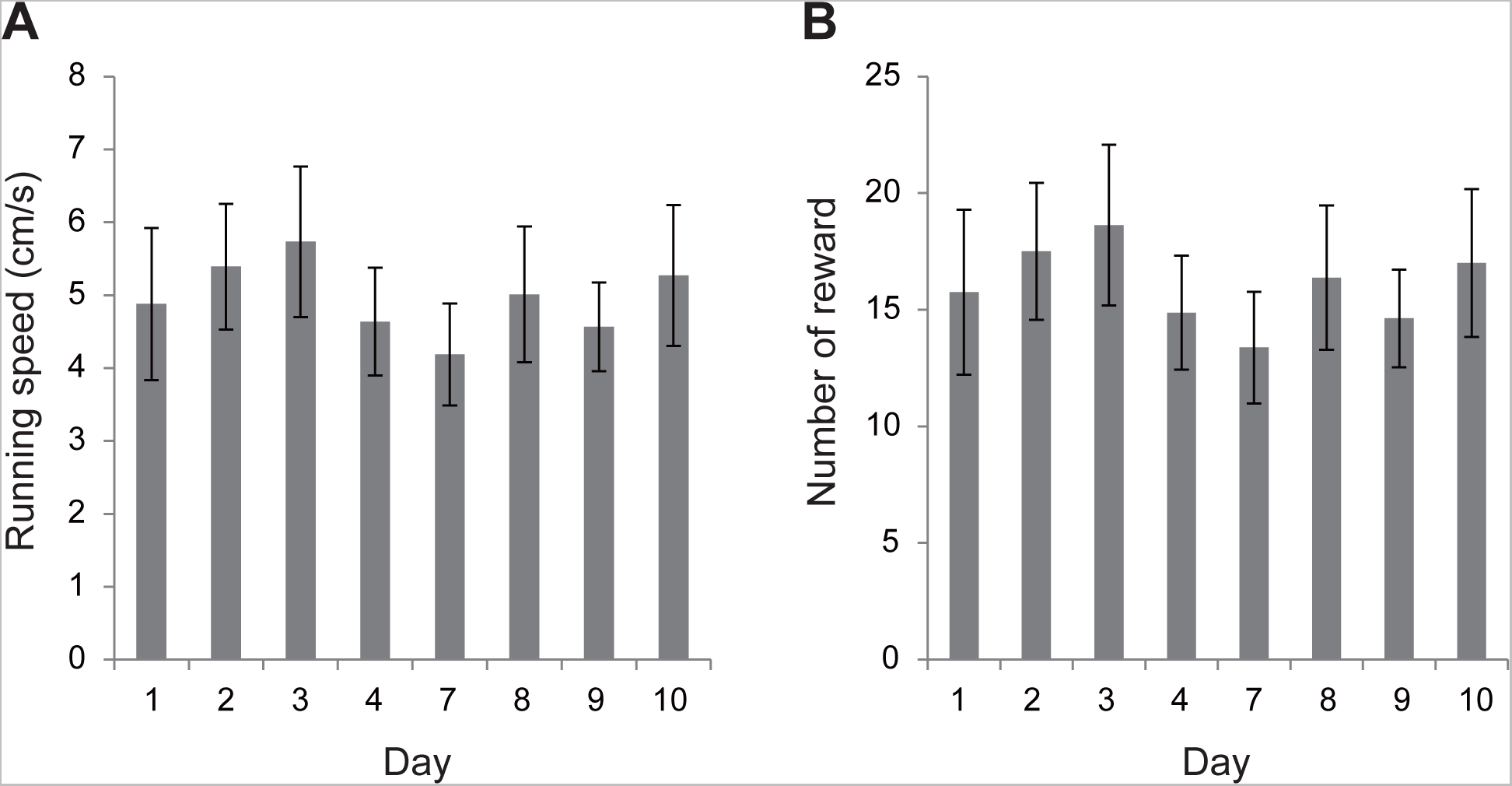
Running speed and reward rate of animals in the long-term recording experiment. (A) Running speed at each recording session. Significance was evaluated by Friedman test (*P* = 0.46, n = 8). (B) Number of rewards at each session. Significance was evaluated by Friedman test (*P* = 0.44, n = 8). Data are shown as mean ± SEM.

**Fig. S2.**
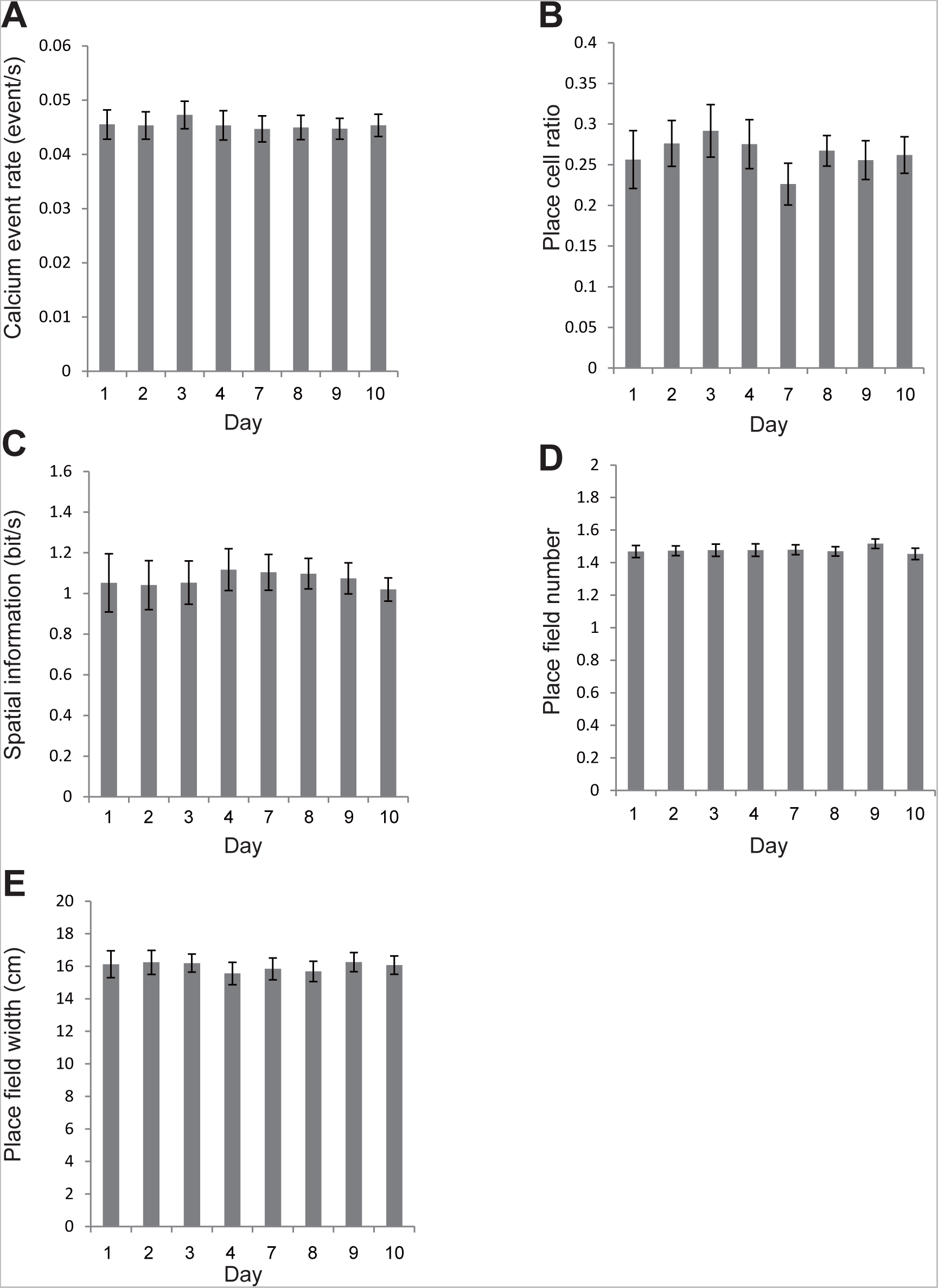
Place cell properties in the long-term recording experiment. (A) Calcium event rate. Significance was evaluated by one-way repeated measures ANOVA (F_2,21_ = 1.5, *P* = 0.19, n = 8). (B) Fraction of cells with significant place cells. Significance was evaluated by Friedman test (*P* = 0.19, n = 8). (C) Spatial information content of place cells per unit time. Significance was evaluated by one-way repeated measures ANOVA (F_2,21_ = 0.33, *P* = 0.94, n = 8). (D) Number of place field for each cell. Significance was evaluated by Friedman test (*P* = 0.79, n = 8). (E) Place field size for each cell. Significance was evaluated by one-way repeated measures ANOVA (F_2,21_ = 0.65, *P* = 0.71, n = 8). Data are shown as mean ± SEM.

**Fig. S3.**
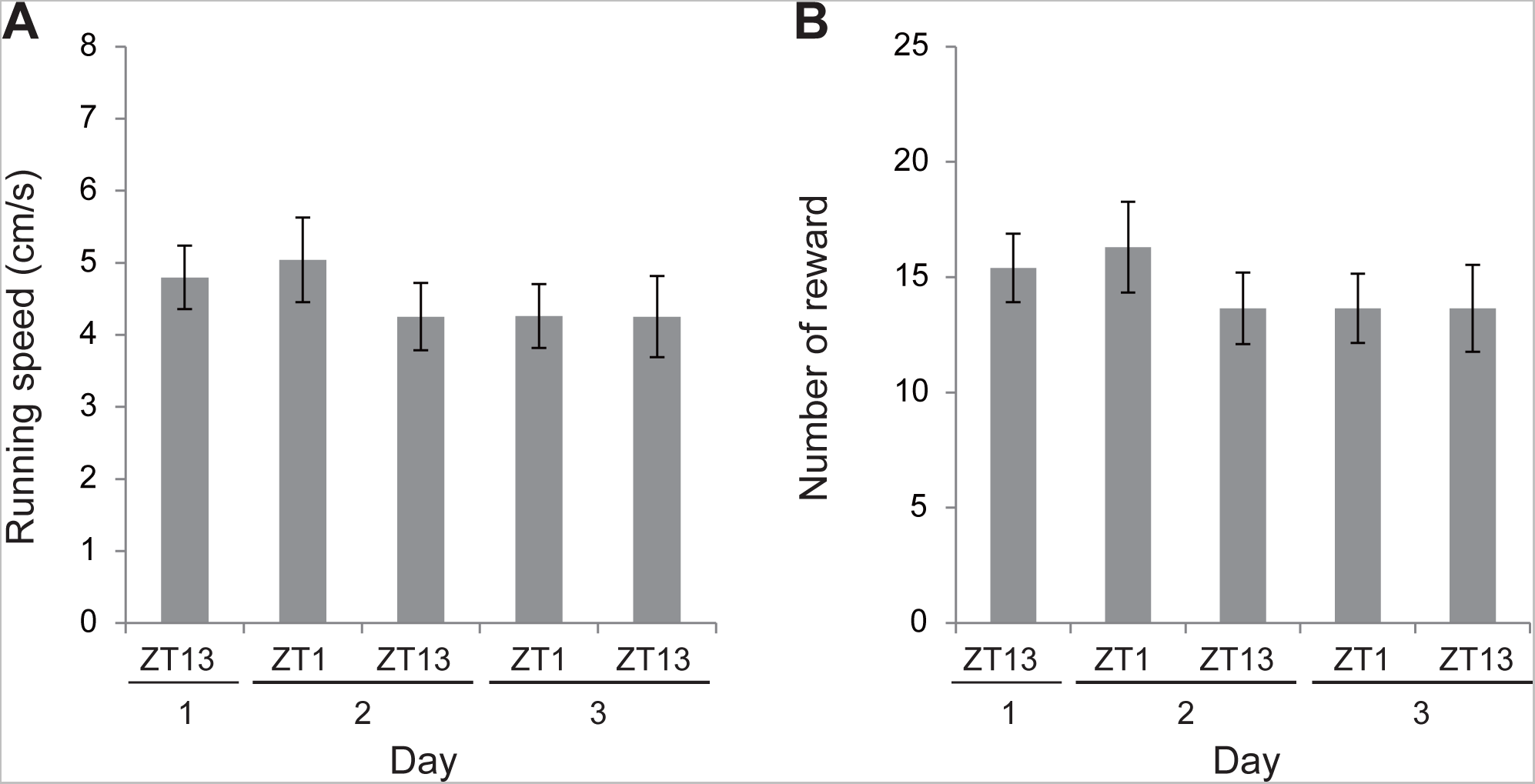
Running speed and reward rate of animals in the bi-daily recording experiment. (A) Running speed at each recording session. Significance was evaluated by Friedman test (*Q* = 6.51, *P* = 0.16, n = 20). (B) Number of rewards at each session. Significance was evaluated by Friedman test (*Q* = 5.53, *P* = 0.24, n = 20). Data are shown as mean ± SEM.

**Fig. S4.**
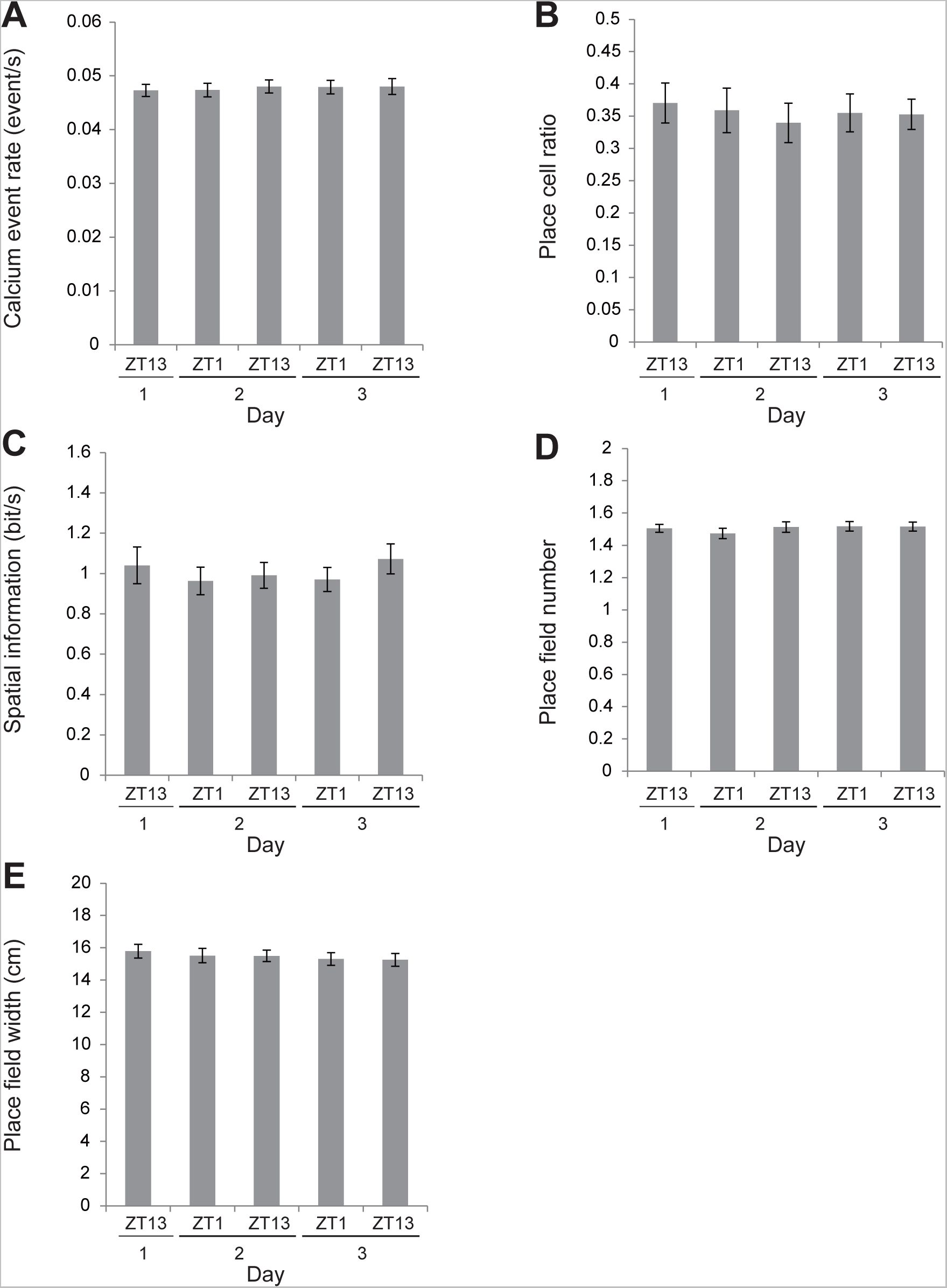
Place cell properties in the bi-daily recording experiment. (A) Calcium event rate. Significance was evaluated by one-way repeated measures ANOVA (F_4,76_ = 0.51, *P* = 0.73, n = 20). (B) Fraction of cells with significant place cells. Significance was evaluated by one-way repeated measures ANOVA (F_4,76_ = 0.45, *P* = 0.78, n = 20). (C) Spatial information content of place cells per unit time. Significance was evaluated by one-way repeated measures ANOVA (F_4,76_ = 1.30, *P* = 0.28, n = 20). (D) Number of place field for each cell. Significance was evaluated by one-way repeated measures ANOVA (F_4,76_ = 0.95, *P* = 0.44, n = 20). (E) Place field size for each cell. Significance was evaluated by Friedman test (*Q* = 5.24, *P* = 0.26, n = 20). Data are shown as mean ± SEM.

**Fig. S5.**
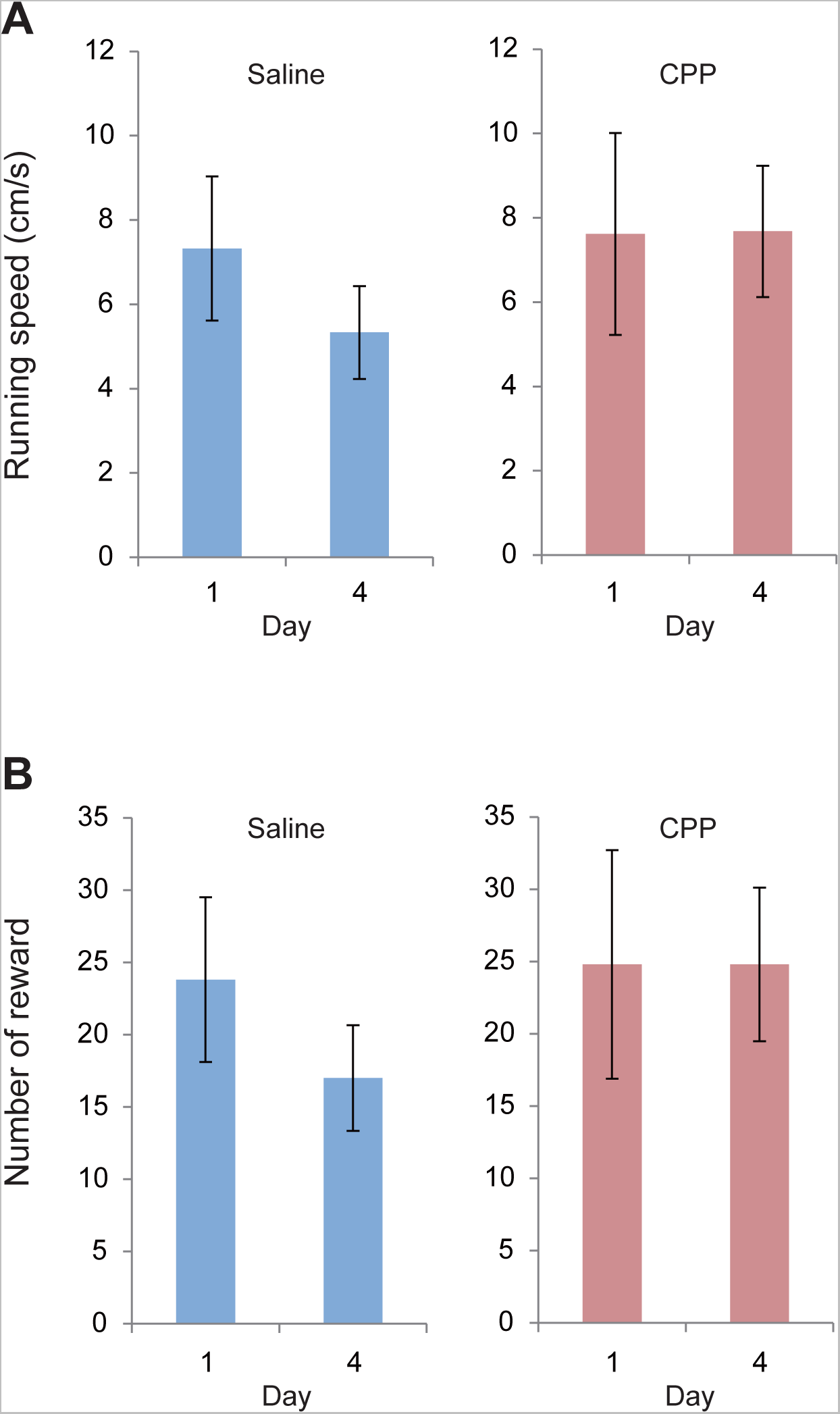
Running speed and reward rate of animals in the CPP experiment. (A) Running speed at each recording session. Significance was evaluated by Wilcoxon signed-rank test (*P* = 0.063, n = 5 for saline; *P* = 1.0, n = 5 for CPP). (B) Number of rewards for each session. Significance was evaluated by Wilcoxon signed-rank test (*P* = 0.063, n = 5 for saline; *P* = 1.0, n = 5 for CPP). Data are shown as mean ± SEM.

**Fig. S6.**
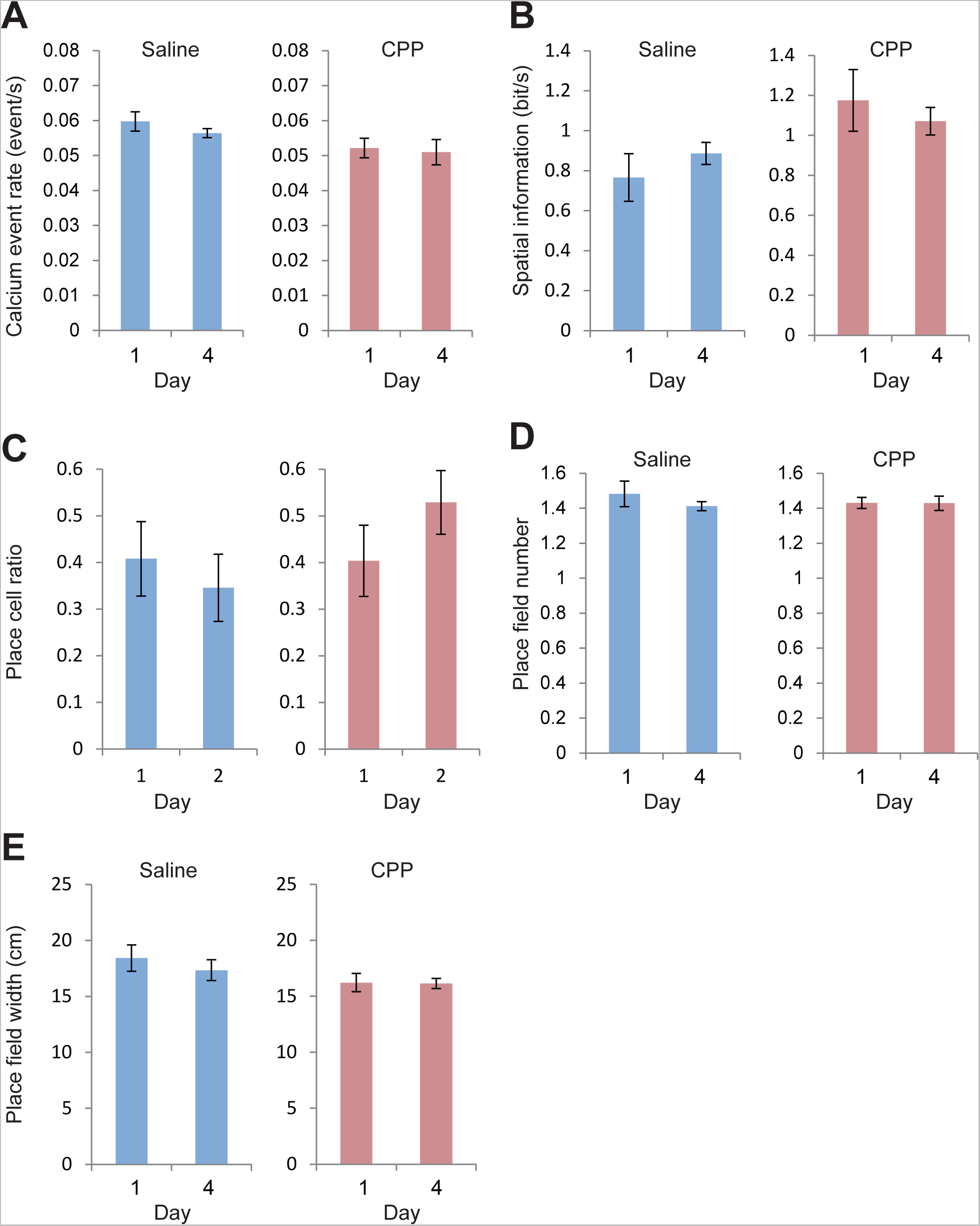
Place cell properties in the CPP experiment. (A) Calcium event rate. Significance was evaluated by Wilcoxon signed-rank test (*P* = 0.31, n = 5 for saline; *P* = 0.44, n = 5 for CPP). (B) Fraction of cells with significant place cells. Significance was evaluated by Wilcoxon signed-rank test (*P* = 0.063, n = 5 for saline; *P* = 0.063, n = 5 for CPP). (C) Spatial information content of place cells per unit time. Significance was evaluated by Wilcoxon signed-rank test (*P* = 0.19, n = 5 for saline; *P* = 0.44, n = 5 for CPP). (D) Number of place fields for each cell. Significance was evaluated by Wilcoxon signed-rank test (*P* = 0.44, n = 5 for saline; *P* = 0.81, n = 5 for CPP). (E) Place field size for each cell. Significance was evaluated by Wilcoxon signed-rank test (*P* = 0.32, n = 5 for saline; *P* = 1.0, n = 5 for CPP). Data are shown as mean ± SEM.

